# Rapid Screening and Validation of Intracellularly Active Antimycobacterial Compounds with Efficacy in Cellular and Murine Tuberculosis Models

**DOI:** 10.1101/2025.09.23.677984

**Authors:** Mahima Madan, Kanika Bisht, Neetu Sehrawat, Vivek Rao

## Abstract

The discovery of effective intracellular antimycobacterial agents is limited by the slow growth of *Mycobacterium tuberculosis* (Mtb) and a lack of scalable models that effectively replicate host-pathogen interactions. To address this, we established a robust *ex vivo* infection model using human monocyte-derived THP-1 macrophages and the fast-growing pathogenic surrogate *Mycobacterium marinum* (Mmar) which recapitulates key features of Mtb intracellular infection, including macrophage colonization and innate immune activation. This model supports medium-throughput screening workflows for assessing antibiotic-mediated control of intracellular mycobacteria. Using this platform, we screened FDA-approved antipsychotics and identified the phenothiazines- trifluoperazine (TFP) and fluphenazine (FFP), as potent adjuncts to frontline anti-tuberculosis therapy (ATT). Combinations of either phenothiazine with frontline TB drugs enhanced intracellular bacterial clearance more effectively than the previously reported antidepressant sertraline, initiating significant control within 12 hours of treatment in macrophages. In murine models of TB infection, TFP and FFP further potentiated bacterial clearance and tissue recovery when used in combination with standard anti-TB therapy (ATT). Together, our results validate the THP1–Mmar infection model as a rapid, robust, and scalable platform for identifying intracellular antimycobacterial agents and host-directed therapeutics, offering a valuable tool to accelerate TB drug discovery and development.

## Introduction

Tuberculosis (TB), despite continued efforts at control, has remained a major global public health problem draining global economies^1^. Identification of novel therapeutic regimens that can effectively shorten the duration of anti-TB therapy (ATT) without contributing to antimicrobial resistance (AMR), is an absolute necessity for a country like India that accounts for more than a quarter of annual infections^2 3^. Recent research efforts have focused on host-directed therapy (HDT) approaches aimed at strengthening innate immune responses of the host to improve disease outcomes and treatment efficacy^4 5 6 7^. In line with this objective, we have previously identified and demonstrated the benefit of an FDA-approved antidepressant-sertraline (SRT) as an adjunct regimen that has shown significant promise in preclinical models of Mtb infection^8 9^. Notably, the SRT-ATT combination has significantly promoted early bacterial clearance, enhanced tissue recovery, and host survival by modulating host innate responses^10^. Importantly, this combination was effective even under conditions of antibiotic insufficiency highlighting the potential of host-augmenting regimens in overcoming current therapeutic limitations^8^. While Mtb-macrophage infection models have been routinely used to evaluate and identify antimicrobial compounds and host-directed therapies, the slow growth rate of Mtb presents a major bottleneck for rapid screening, lead prioritization, and preclinical development of therapeutic candidates^11 12 13 14^. To address this, the fast-growing non-pathogenic *Mycobacterium smegmatis* (Msm) has frequently been used as a surrogate in the early phases of high-throughput screening efforts^15^. However, substantial differences in infection dynamics and host immune responses between Msm and Mtb limit the utility of Msm-based models, particularly for identifying compounds that act through modulation of macrophage-mediated innate immune pathways^16 17 18^.

To overcome this lacuna, we have developed an *ex vivo* infection model with Mmar that recapitulates the intracellular growth kinetics of Mtb in THP1 macrophages. Having established the dose-response kinetics of frontline TB drugs, we employed this model as a screening platform to identify putative antidepressants and antipsychotics that could enhance antibiotic efficacy. We identify two phenothiazines - trifluoperazine (TFP) and fluphenazine (FFP) as adjuncts that significantly supplement the bacterial clearance by frontline TB drugs in macrophages. We further demonstrate that the combinations with TFP or FFP significantly accelerate bacterial clearance in murine models of infection. These findings highlight the convenience of the Mmar–THP1 infection model as a rapid, scalable, and biologically relevant screening platform for time-efficient lead discovery and validation of novel intracellular antimycobacterial agents and HDTs, thereby contributing to the ongoing efforts to develop effective TB therapies.

### Methodology Cell Culture

THP1 (ECACC) and THP1 dual monocytes ((thpd-nfis, InvivoGen, France) were cultured in RPMI 1640 containing 10% FBS and 1 mM sodium pyruvate at 37^0^C in a 5% CO_2_ humidified incubator. They were differentiated into macrophages using 100 nM Phorbol-13-myristate 13-acetate (PMA) for 24 hours, followed by a resting period of 48 hours, before use for the experiments.

### Bacterial Strains and Media

Mtb Erdmann and Mmar were grown in Middlebrook 7H9 medium with 4% Albumin-dextrose-saline with 0.05% Tween 80 for liquid cultures and on Middlebrook 7H10 plates supplemented with 10% oleic acid-albumin-dextrose-catalase (OADC) at 37^0^C.

### Reagents

HiglutaXL RPMI-1640, fetal bovine serum (FBS) and Rifampicin (CMS1889) and were purchased from Himedia laboratories, India. PMA (P8139), Isoniazid (I3377), Pyrazinamide carboxamide (P7136), Ethambutol dihydrochloride (E4630), Sertraline hydrochloride (S6319), Trifluoperazine hydrochloride (T6062), Fluphenazine hydrochloride (PHR1792), Chlorprothixene hydrochloride (C1671), Amikacin sulfate (A2324) were purchased from Sigma-Aldrich, USA.

### Macrophage infection and Intracellular growth assays

Log phase mycobacterial cultures were centrifuged and washed with PBST (1x PBS with 0.05% Tween 80) twice, and a single cell suspension was made after spinning down the culture at 150g for 10 minutes. THP1 dual macrophages were infected (Mtb for 6 h or Mmar for 3 h) at MOI5. The cells were then washed twice with 1X PBS, treated with amikacin for 1 h to remove extracellular bacteria and incubated. At specific time points in the assay, cells were lysed in water containing 0.05% Tween 80 and plated as serial dilutions on 7H10 agar plates and incubated at 30^0^C for a week (for Mmar) or at 37^0^C for 3-5 weeks (for Mtb).

### Imaging experiments

For imaging using mCherry-tagged Mmar, seeding of THP1 macrophages was done in 96-well treated plates at a density of 0.1 million cells per well and infected with Mmar tagged with mCherry at an MOI of 5 for 1 hour. Cells were treated with antibiotics and, thereafter, at specific time points, cells were imaged in a Floid imaging station (from ThermoFisher Scientific).

### Mice infection and antibiotic treatment

6 to 10 weeks old C3HeB/FeJ or BALB/c mice were infected with 500 CFU of Mtb via the aerosol route in a Glascol inhalation chamber. After 4 weeks (for BALB/c mice) or 2 weeks (for C3HeB/FeJ mice) of infection, animals were given antibiotics or drugs of interest *ad libitum* in the drinking water for 8 weeks (twice a week). Drugs were used in the following concentrations-H (100 mg/L), R (40 mg/L), Z (150 mg/L), E (100 mg/L), and TFP/ FFP (3 mg/l)^19 20^. For survival assessment, 10 C3HeB/FeJ animals per group were monitored regularly. For assessing bacterial counts in BALB/c mice, animals were euthanized at 4-, 6-, and 9-weeks post-treatment. The lungs and spleen of infected animals were collected in normal saline (sterile), homogenized and plated on 7H11 agar plates containing OADC and incubated at 37^0^C for estimating bacterial numbers (CFU). The left caudal lung lobe was fixed in 4 % formaldehyde, sectioned and stained with H & E for histopathology studies.

### Luminescence Assay

THP1 dual cells stably expressing Lucia luciferase were seeded at a density of 60,000 cells per well in a 96-well plate and infected at MOI5. Cells were either left untreated or treated with appropriate drug combinations, and supernatant was collected after incubation for specific time points. The extent of luminescence in the supernatant was estimated using QUANTI-Luc (InvivoGen, France) detection reagent as per the manufacturer’s recommendations.

### Statistical analysis

The results are depicted as mean ± SEM. Statistical analysis was carried out using Student’s t-test (Welch’s test or Mann-Whitney test) using GraphPad Prism. P values less than 0.05, 0.01, and 0.005 are represented as *, **, and ***, respectively.

## Results

### Development of a THP1-Mmar infection model that mimics the infection dynamics of Mtb

In order to develop a fast and robust screen for intracellular control of mycobacterial infection, we tested the ability of Mmar to infect THP1 monocyte-derived macrophages at different infection densities and assessed the growth of bacteria (Figure 1A). The kinetics of infection mirrored the initial inoculum dose with Mmar at an MOI of 0.1 showing significantly lower bacterial numbers than the other doses at all time points (Figure 1B). While the higher doses of 1, 5, and 10 also showed a dose-dependent increase in bacterial numbers, the growth rates relative to the initial uptake were comparable across 96h of infection. In fact, a gradual increase in intracellular bacterial numbers was observed with time, irrespective of the dose of infection. Interestingly, infection with an MOI of 10 induced significant lysis of macrophages by the 96h time point, similar to the high levels of necrotic cell death observed in a high-dose Mtb infection model of THP1 macrophages.

**Figure 1:**
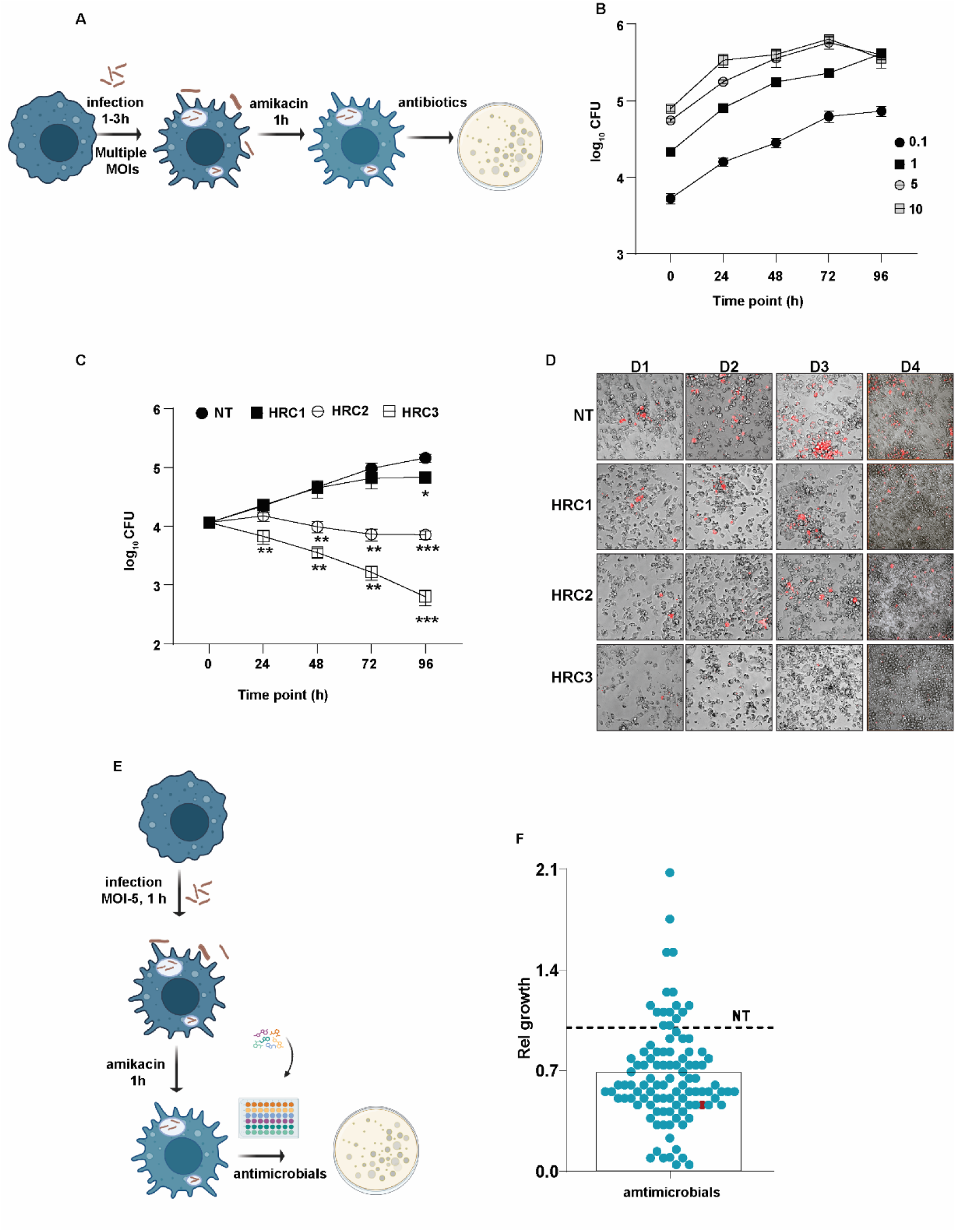
Establishment of a robust THP1-Mmar infection model for rapid screening of antimycobacterials. A) Workflow depicting the Mmar infection strategy for THP1 macrophages. B, C). Growth kinetics of Mmar in THP1 macrophages following infection at different MOIs (0.1, 1, 5, 10) (B) or (C) in the presence of different concentrations of the frontline TB drugs-isoniazid (H) or rifampicin (R) - HRC1-H=2 ng/ml, R=10 ng/ml; HRC2-H=20 ng/ml, R=100ng/ml; HRC3-H=200 ng/ml, R=1000 ng/ml. The bacterial numbers (CFU) were calculated at indicated time intervals post-treatment and are represented as mean ± SEM of triplicate assays from 3 independent experiments. D). Infection kinetics of mCherry-tagged Mmar in THP1 macrophages, left untreated (NT) or treated with different concentrations of HR. At indicated time intervals, cells were imaged using the FLoid Cell imaging station. A representative image is depicted. E, F). The schematic used in this study for the use of this model as a platform for rapid screening is shown in E. Previously identified antimicrobials were evaluated for their ability to limit intracellular growth of Mmar in this model and is depicted as relative CFU in the test samples with respect to untreated controls (NT, dotted line) in triplicate assays. The CFU numbers in HR treated samples is shown in red.

The combination of the first-line TB drugs isoniazid (H) and rifampicin (R) was extremely effective in controlling infection in a dose and time-dependent manner (Figure 1C). A dose of H=2 ng/ml, R=10 ng/ml (HRC1) failed to control bacterial growth at any time point in the assay. A 10-fold higher dose (HRC2) was relatively bacteriostatic until 96 h of treatment. However, this dose reduced bacterial numbers by 4-5-fold by 48 h of treatment in comparison to untreated cells, despite failing to impact growth at 24 hours of treatment. In contrast, the dose of HR: H=200 ng/ml, R=1000 ng/ml (HRC3) restricted infection by 4-5-fold even by 24 h that escalated to a 70-fold lower bacterial numbers by 96h of treatment. This pattern of infection and control was also reflected in cells infected with a fluorescently labeled mCherry-tagged Mmar (Figure 1D). While untreated cells showed extensive growth by day 4, cells treated with escalating doses of HR showed a progressive decrease in bacterial numbers with antibiotic dose and time of treatment.

To evaluate this infection model as a platform to screen for antimycobacterials, Mmar-infected THP1 macrophages were treated with 143 antimicrobial compounds from the Spectrum collection (MicroSource Discovery Systems) library in a blinded screen, and the growth was monitored at 48 h following treatment (Figure 1E). Out of the ∼65 molecules that controlled intracellular Mmar growth, more than 55 compounds were previously identified to possess antimycobacterial properties, thereby establishing the veracity and robustness of the infection model as a platform for lead molecules with mycobactericidal

### THP1-Mmar infection is effective to screens adjunct therapy regimens

We have previously identified a potent adjunct therapy regimen of ATT and sertraline that complements the frontline TB drugs in enhancing Mtb kill kinetics ^8^. To assess if the current infection model would provide a comprehensive read out for host targeting therapies, we tested the effect of adding SRT to HR on Mmar growth. Again, infection control dynamics with fluorescent bacteria (Figure 2A) mirrored the analysis by CFU-based growth assays (Figure 2B, C). Even at the dose of HRC1 that showed negligible impact, addition of SRT significantly reduced bacterial numbers up to 125-fold by day 4 treatment (Figure 2B). While treatment with HRC2 alone reduced bacterial numbers significantly, addition of SRT further decreased bacterial numbers by 70-fold by day 4 of treatment, advocating the usefulness of this model as a screen for novel host-directed adjunct modalities (Figure 2C).

**Figure 2.**
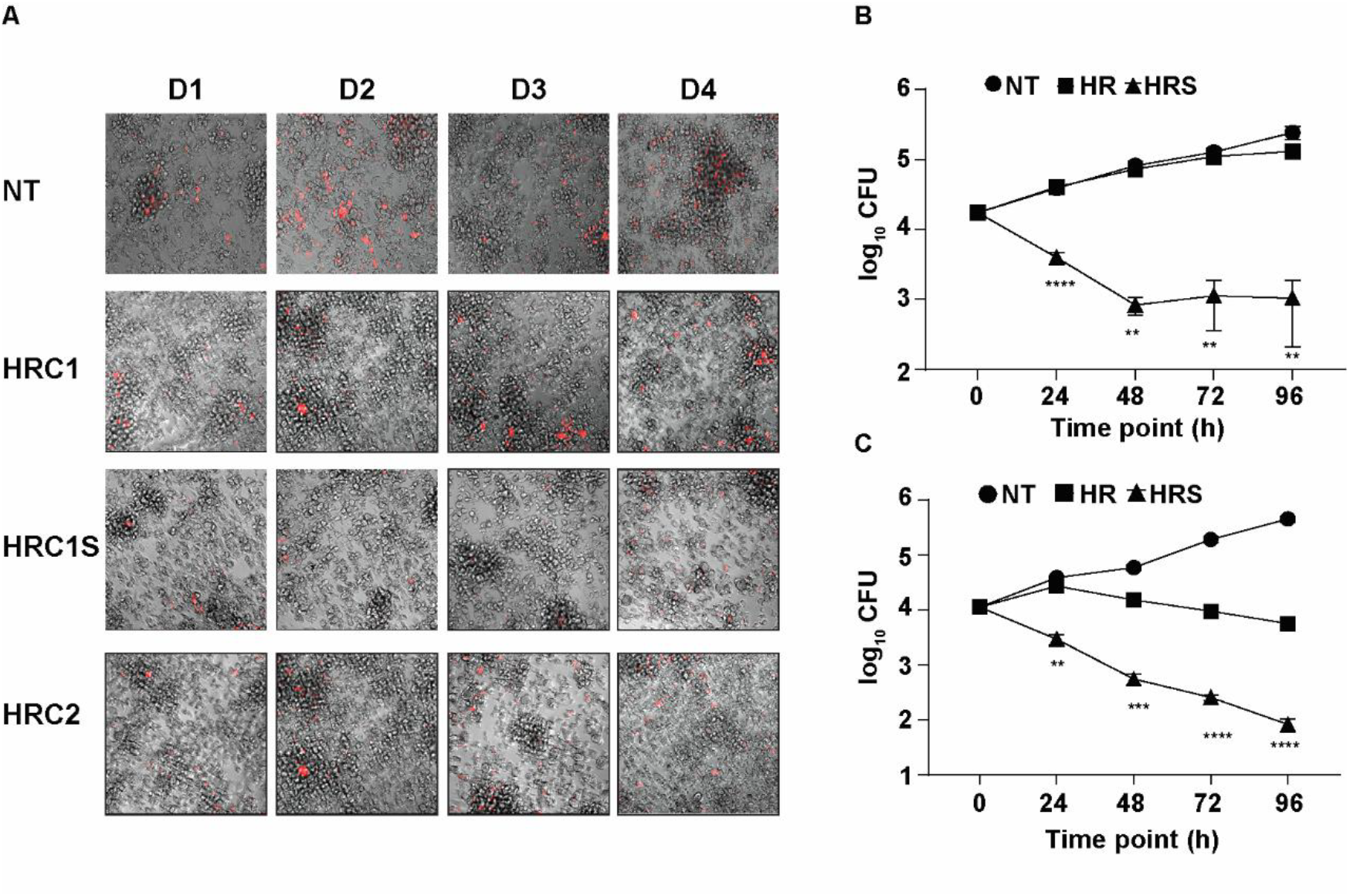
-THP1-Mmar infection model reflects the enhanced effects of potential HDTs. A-C) Infection kinetics of mCherry tagged Mmar in THP1 macrophages that were either left untreated (NT) or treated with different concentrations of HR (HRC1-H=2 ng/ml, R=10 ng/ml; HRC2-H=20 ng/ml, R =100ng/ml) alone and in combination with SRT at 20 µM (HRS). At various time intervals, the cells were imaged using FLoid Cell imaging station (A). A representative image is depicted. The bacterial numbers (CFU) were calculated at different time intervals post treatment (24, 48, 72, 96 h) and are represented as mean± SEM from 3 independent experiments (B, C).

### Screen identifies other antipsychotics and antidepressants (ADAPs) as more effective HDTs

Given our identification of SRT (an anti-depressant) as a powerful adjunct to ATT, we explored if other antidepressants and related antipsychotics could enhance bacterial control in combination with TB drugs. We sourced 76 designated ADAPs from the Spectrum collection chemical library and tested their ATT adjunct properties in the THP1-Mmar infection model (Figure 3A). While most of the compounds failed to modify growth restriction by HR, 13 compounds significantly reduced the bacterial burdens by >1.8-fold relative to HR alone by 48h of treatment (Figure 3B, C). Further analysis of these compounds against Mtb infections revealed distinct groups in the ability of the compounds to abet antibiotic efficacies (Figure 3D). While venlafaxine, desvenlafaxine, citalopram, desipramine did not affect the growth control efficiency of HR, protriptyline, amitriptyline, chlorpromazine and duloxetine showed similar efficiency as SRT in enhancing HR dependent Mtb growth restriction. However, two of the phenothiazines-trifluoperazine (TFP) and fluphenazine (FFP) and chlorprothixene significantly enhanced Mtb growth control by more than 5-6-fold in comparison to the effect of HRS mediated bacterial control (Figure 3D).

**Figure 3:**
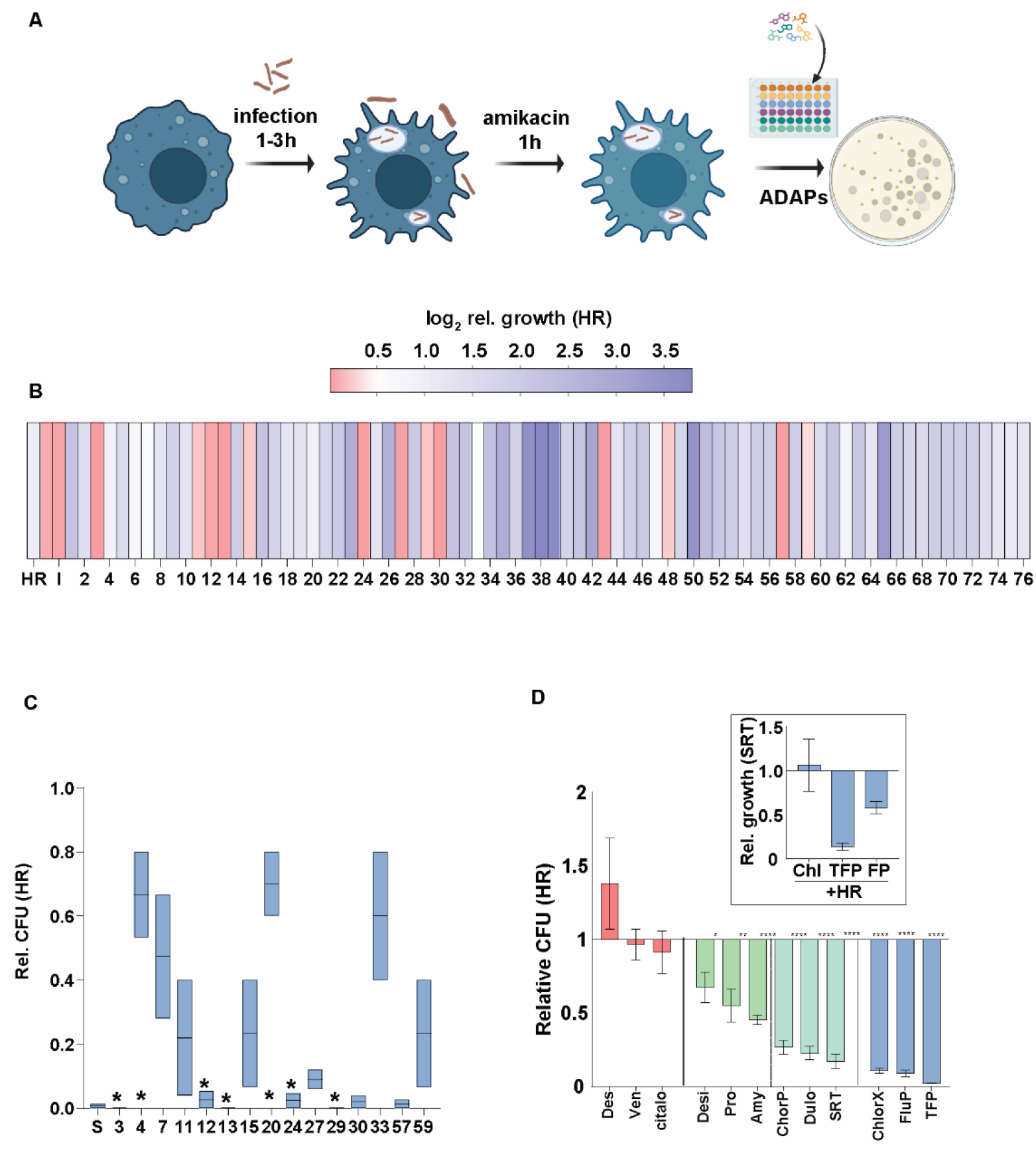
TFP and FFP augment Mtb control in combination with frontline TB drugs. A-D). The schematic for testing FDA approved ADAPs as antibiotic adjuncts (A). The intracellular growth of Mmar in THP1 macrophages after 24 h of treatment with the 76 previously identified ADAPs is depicted as heatmap of average CFU numbers in the test (HR + ADAP) samples relative to the control HR alone samples from 2 independent experiments (B). The intracellular growth of Mmar (C) or Mtb (D) in macrophages that were left alone (NT) or treated with HR alone and in combination with SRT (S) or selected ADAPs from the screen. Data represents mean relative CFU (HR+ ADAP/HR) + SEM of triplicate assays from 3 independent experiments (N=3). Inset in panel D) depicts the comparative CFU in HR + ChlorX, HR+TFP or HR+FFP treated samples relative to HRS treated samples as mean CFU ± SEM of triplicate assays from 3 independent experiments (N=3).

To further investigate the efficacy of the selected phenothiazines as a potential HDT, the effect of TFP and FFP on intracellular Mtb growth was analyzed temporally. In contrast to the negligible impact of SRT to enhance HR dependent growth control at 24 h of treatment, both TFP and FFP decreased bacterial numbers as early as 12 h after treatment by 4-fold and 2-fold relative to HR alone, respectively (Figure 4A), This reduction further increased to around 10-15-fold by day 3 and approximately 200-fold by day 5 of treatment (Figure 4B, C). More importantly, the effect of these molecules followed dose-dependent kinetics with a significant decrease (relative to HR) at 5µM of either molecule after 72 h, and doses as low as 1µM showed a significant effect by day 5 of treatment.

**Figure 4:**
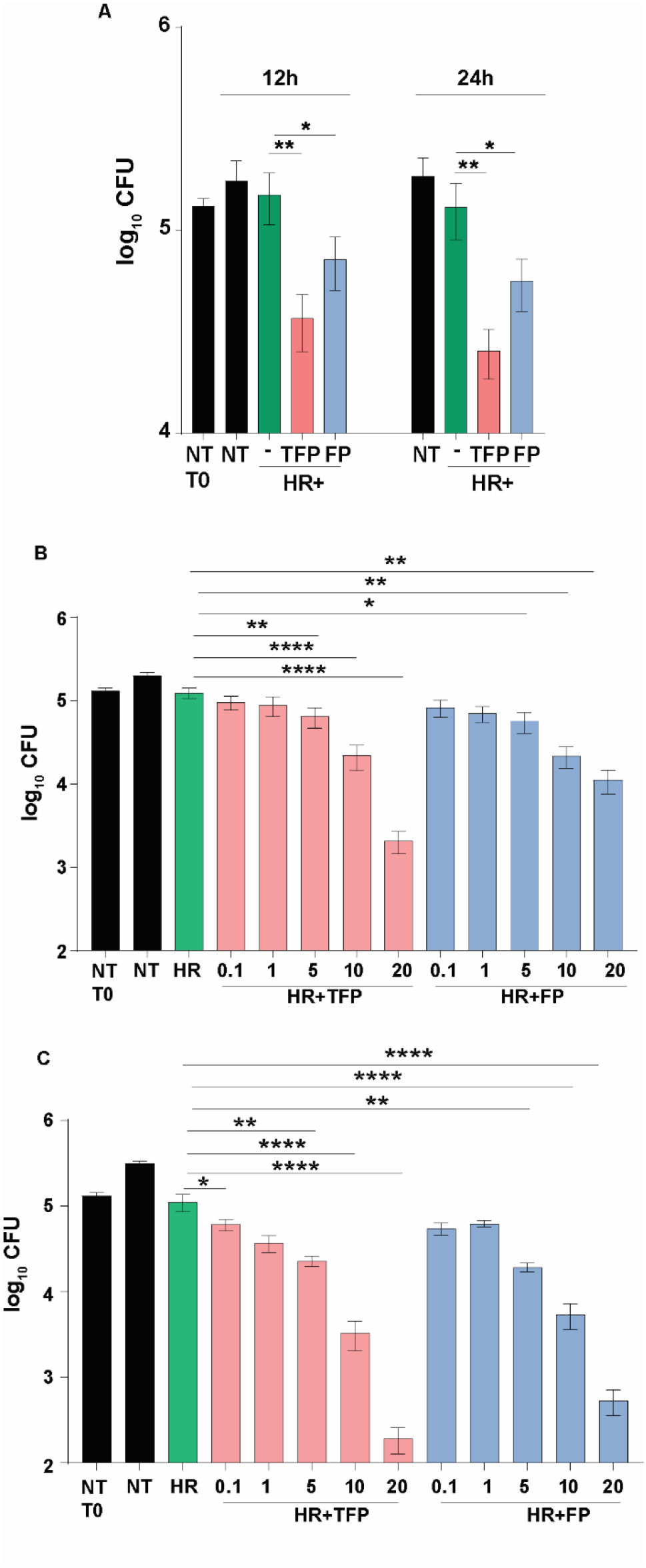
TFP and FFP impart bacterial early control in macrophages. A). The Intracellular bacterial numbers in Mtb-infected THP1 macrophages left untreated (NT) or treated with HR alone or in combination with 20 µM of TFP or FFP for 12 h and 24 h The bacterial numbers are represented as mean CFU ± SEM from 3 independent experiments (N=3). B-C) Mtb growth in THP1 macrophages left untreated (NT) or treated with HR alone or in combination with different concentrations (0.1 µM,1 µM, 5 µM,10 µM, 20 µM) of TFP and FFP at day 3 (B) and day 5 (C) post-infection is represented as mean CFU ± SEM from 3 independent experiments (N=3).

### Both the phenothiazines effectively restrict Mtb induced type I IFN responses in macrophages

Given our use of SRT as a HDT for TB based on its tested ability to restrict infection induced type I IFN response, we investigated if TFP and FFP could also inhibit this response in macrophages. Indeed, following infection with Mmar, THP1 macrophages elaborated type I IFN in an infection dose and kinetic manner (Figure 5A). The levels were significantly higher in the infected samples by 6h with a steady increase till 48h of infection, the higher MOI of 5 consistently induced significantly higher type I IFN response levels than the lower MOI of 1. Both TFP and FFP inhibited Mtb induced type I IFN response in THP1 macrophages, corroborating the correlation between type I IFN inhibition and the ability of compounds to control Mtb growth (Figure 5B).

**Figure 5:**
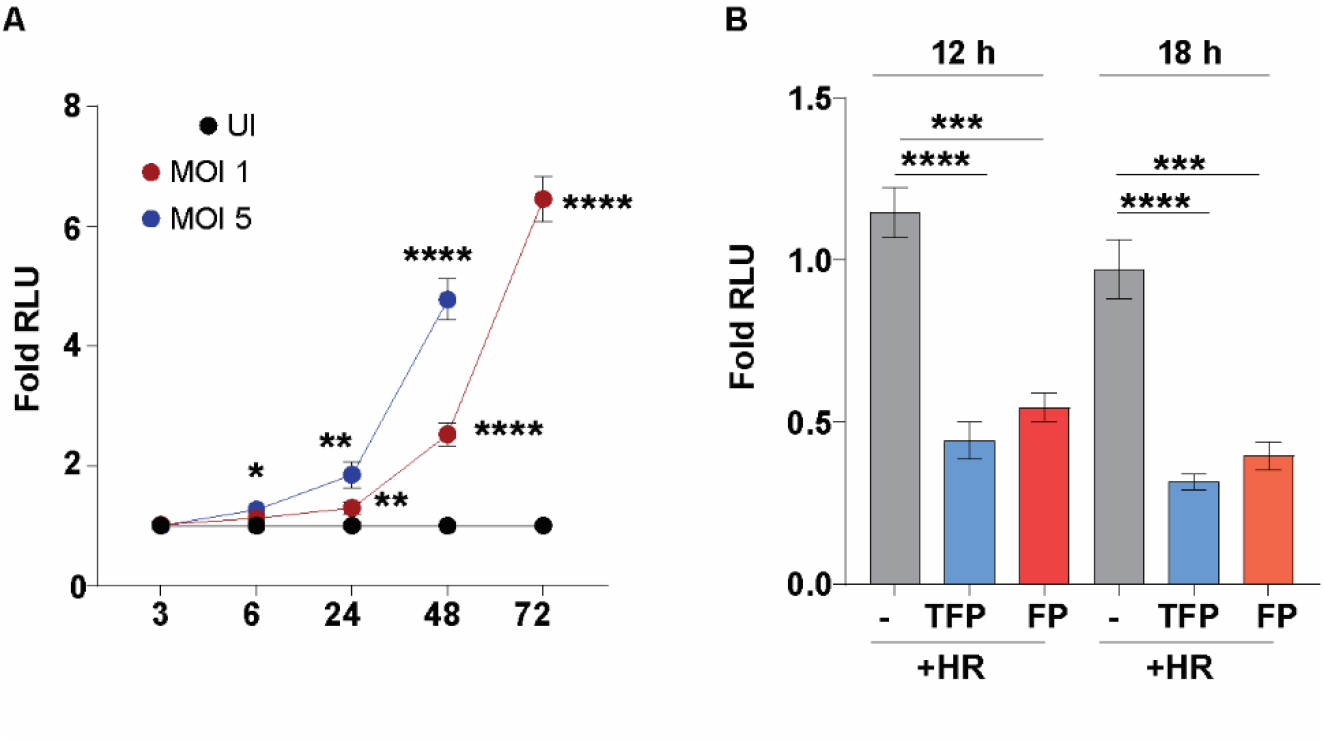
Phenothiazines restricts Mtb induced type I IFN responses. A) The induction of type I IFN in uninfected THP1 dual macrophages or after infection with two different MOIs of Mmar (1 and 5) was evaluated at 3, 6, 24, 48, and 72 h post-infection. The IRF-dependent luciferase activity is represented as mean RLU ± SEM of triplicate assays from 3 independent experiments (N=3). B) The IRF-dependent luciferase activity in Mtb infected THP1 dual macrophages left untreated or treated with HR alone for or in combination with 20 µM of TFP or FFP for 12 h and 24 h is depicted as mean RLU ± SEM of triplicate assays from 3 independent experiments (N=3).

### Enhanced bacterial control by the combination of TFP or FFP with frontline TB drugs in the murine model of infection

Both TFP and FFP were evaluated in the previously documented acute and chronic infection models of murine TB treated with suboptimal concentrations of antibiotics to allow for efficient assay of antibiotic boosting^8^ (Figure 6A). The highly sensitive C3HeB/FeJ mice succumbed to infection by ∼100 days on infection with a dose of 500 CFU of Mtb. While the treatment with HR at the suboptimal concentration did not lend any benefit to host survival, the addition of either TFP or FFP significantly delayed the host kill kinetics, with all animals succumbing to infection by ∼200 days of infection (Figure 6B). Addition of TFP or FFP to HRZE significantly reduced CFU (4-5-fold lower) relative to HRZE alone in the lungs of BALB/c mice by 4 and 6 weeks of treatment (Figure 6C). The increase in control with the combination was not reflected after 9 weeks of treatment, in part due to the heightened control of bacterial numbers by HRZE alone (Figure 6C). However, despite insignificant differences in the extent of lung lesions of the HRZE and HRZETFP/ HRZEFFP treated animals at 4 and 6 weeks, there was a marked decrease in the lesions of mice treated with HRZETFP in comparison to HRZE alone by 9 weeks of treatment (Figure 6D) further validating the multiple levels of advantage afforded by the combination of frontline TB drugs and TFP/ FFP. Taken together, this work highlights the potential of this robust screen in identifying novel antimycobacterials with translational significance, allowing for the fast-track movement of novel entities towards preclinical/ clinical trials.

**Figure 6:**
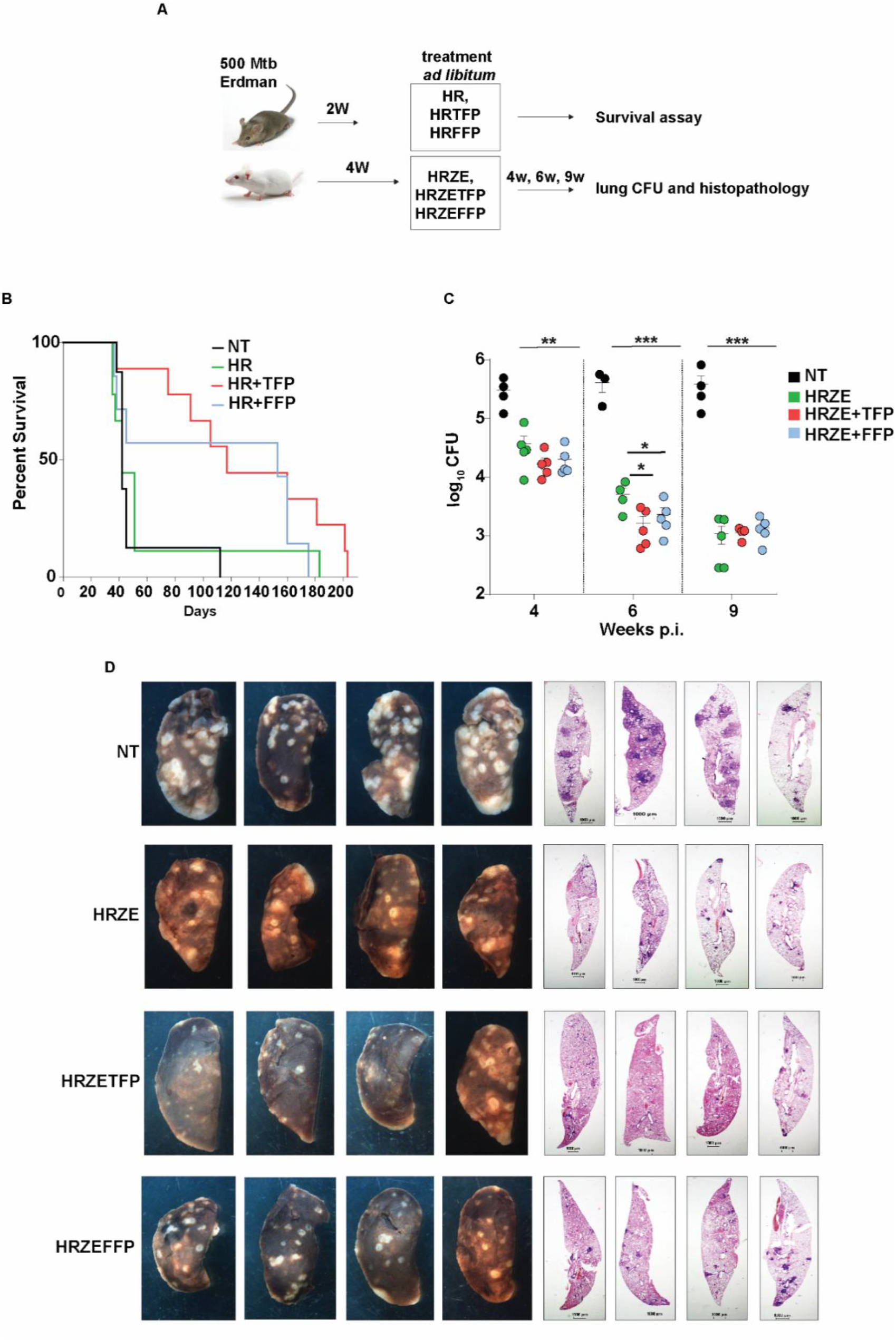
TFP and FFP control Mtb growth in combination with ATT *in vivo*. A) Schematic for the *in vivo* testing of TFP and FFP in the murine model of infection. The drugs were given *ad libitum* in drinking water after 2 or 4 weeks of infection with 500 CFU of Mtb. B) Survival of Mtb-infected C3HeB/FeJ mice left untreated (NT) or treated with HR (H=1 mg/L and R=0.4 mg/L) alone or in combination with TFP (3 mg/L) or FFP (3 mg/L) is depicted in a Kaplan-Meier survival analysis curve with 10 animals in each group. C) Bacterial numbers in the lungs of Mtb infected Balb/c mice either left untreated or treated for 4 weeks, 6 weeks and 9 weeks with HRZE alone (H=100 mg/L, R=40 mg/L, Z=150 mg/L, E= 100 mg/L) or in combination with TFP (3 mg/l) or FFP (3 mg/l) are depicted as mean CFU± SEM from 4-5 mice per group. D) The gross morphology of lungs of Mtb infected Balb/c mice infected left untreated (NT) or treated with HRZE (H=100 mg/l, R=40 mg/l, Z=150 mg/l, E=100 mg/l) alone or in combination with TFP (3 mg/l) or FFP (3 mg/l) for 9 weeks is depicted. The histopathological changes are represented in the H&E-stained lung sections of animals in the different groups.

## Discussion

Development of efficient, robust models for high throughput screens of large numbers of molecules is critical in satisfying the growing need for alternative regimens to accelerate TB treatment. While the use of surrogate mycobacteria like Msm help circumvent the longer assay times dependent on the slow growth rates of Mtb^15, 21^, the lack of coherence in infection patterns associated with two organisms ^16, 22^ restricts its use as a screen for HDTs that rely on host cellular response pathways. In contrast, Mmar combines pathogenicity with a rapid growth profile, enabling shorter assay durations without compromising biological relevance^23^. Our model THP1-Mmar infection successfully recapitulates Mtb-like intracellular replication and responds predictably to frontline drugs, validating its suitability for HDT evaluation. Using this system, we have identified trifluoperazine (TFP) and fluphenazine (FFP), two phenothiazine derivatives, as promising HDTs that exert early, enhanced bactericidal activity compared to sertraline (12 h vs 72 h). This effect corresponds with their capacity to repress infection-induced type I interferon responses in macrophages, reinforcing the pathogen beneficial role of this immune signaling axis.

The *in vivo* adjunctive efficacy of TFP and FFP combined with HR was significant primarily as early as 6 weeks of treatment. This reflects the pharmacokinetic profile of drug delivery via drinking water, with the ability to reach therapeutic concentrations governing the efficacy initiation. We hypothesize that both TFP/FFP accelerate effective HR concentrations, thereby enhancing early bacterial killing, a finding that is supported by our previous observation of sertraline enhancing the PK/PD of TB drugs without affecting clearance in preclinical models^9^. By nine weeks, the comparable bacterial burden between HR alone and combination-treated animals suggests the saturation of antimicrobial effect with prolonged therapy.

Interestingly, despite comparable intracellular activity, TFP was better at the restoration of lung tissue architecture compared to FFP. Given the fact that FFP is used as an injectable clinically, while TFP is given orally, this incongruence could arise from differences in their bioavailability in this study, affecting tissue penetration, or host immune interactions^24 25^. Delineating tissue availability and distribution of different drugs in detail would be necessary to address these issues and would form a part of future studies.

The use of sub-MIC concentrations (0.01x and 0.1x HR) for effective bacterial control with potential HDTs coupled with an intrinsic compatibility of fluorescence- and growth-based readouts, facilitates robust high-throughput imaging assays, improving assay sensitivity and throughput by our model. Overall, the Mmar THP-1 infection model represents a valuable, robust and time-efficient platform for accelerating the discovery of host-directed therapies, thereby aligning well with the global efforts on enhancing TB control through innovative drug development.

## Author contribution

MM, KB, NS and VR were instrumental in the design of the work. MM and KB were involved in the conduct of the work. None of the authors have any conflict of interest.

## Acknowledgements

The authors thank CSIR (VR-MLP2106) for supporting the study. CSIR-STS0016 is acknowledged for continuous maintenance of BSL3 and ABSL2 facilities. The student fellowships from CSIR-India (MM and KB) and UGC-India (NS) are acknowledged.

